# Dynamic Size-Weight Changes After Object Lifting Reduce the Size-Weight Illusion

**DOI:** 10.1101/662999

**Authors:** Vonne van Polanen, Marco Davare

## Abstract

In the size-weight illusion, the smaller object from two equally weighted objects is typically perceived as being heavier. One explanation is that the mismatch between the weight expectation based on object size and actual sensory feedback influences heaviness perception. In most studies, the size of an object is perceived before its weight. We investigated whether size changes would influence weight perception if both would be perceived simultaneously. We used virtual reality to change the size and weight of an object after lifting and asked participants to judge whether objects became lighter or heavier. We found that simultaneous size-weight changes greatly reduced the size-weight illusion to perceptual biases below discrimination thresholds. In a control experiment in which we used a standard size-weight illusion protocol with sequential lifts of small and large objects in the same virtual reality setup, we found a larger, typical perceptual bias. These results show that the size-weight illusion is smaller when size and weight information is perceived simultaneously. This provides support for the prediction mismatch theory explaining the size-weight illusion. Furthermore, these findings suggest that the lifting phase is a critical time window during which brain mechanisms comparing perceived and expected weight mediate the size-weight illusion.

## 1 INTRODUCTION

The size-weight illusion (SWI) is a strong illusion that continues to attract the interest of many researchers since its discovery by Charpentier ^1^. In this illusion, two objects of equal mass but different size are typically perceived to have different weights. Specifically, the smaller object is perceived to be heavier than the larger one. The SWI is the strongest weight illusion ^2^ and can be experienced both when size is perceived haptically or visually ^3^. Despite over a century of research, this illusion remains intriguing and there are still different theories accounting for the SWI.

First, some theories consider a top-down influence. This reasoning involves a mismatch between predicted and perceived weight when lifting the object. Although the mismatch between expected and actual sensory feedback seems important to induce the SWI, previous research suggests that the illusion does not have a purely sensory basis. When initially lifting equally weighted small and large objects, fingertip forces are larger for large compared to small objects ^4^. This suggests that the inappropriate force scaling would influence perceived object weight where the necessary force is less than expected. However, fingertip forces scale to actual object weight after a few lifts whereas the illusion remains ^5^. Besides a purely sensory mismatch, a more cognitive approach assumes that the expectation of object weight based on its size is included into the final weight judgement (see for reviews ^2,6^). This expectation is implicit ^7–9^ and only after considerable practice time the SWI can be extinguished or even reversed ^10^. Because the illusion is shifted in the opposite effect of the expectation, the SWI has sometimes been called anti-Bayesian ^11^.

Secondly, other theories rely on bottom-up explanations, where there is no incorporation of a prediction, but directly available information is used to form a weight percept. For instance, it has been suggested that the SWI is related to the perception of density (i.e. the relation between object size and weight) ^12,13^. It has been shown that density as well as the SWI activate the dorsal premotor cortex, whereas size and weight alone are not represented in this area ^14^.

An issue that has received less attention in the SWI, is the specific timing when this illusion is formed. In most experiments, the object is seen before it is lifted. The weight of an object can be predicted from its size, before it is actually lifted. However, its actual weight can only be veridically perceived after lifting. Therefore, the size of the object is perceived *before* its weight. Moreover, it has been shown that viewed size *before* lifting is enough to induce the illusion^15^ (but see also ^16^ who did not find this). Even if the object is lifted without vision, object size is still haptically perceived earlier than object weight, because size can be perceived through the grasp aperture when fingertips contact the object while weight only apparent after lifting the object. Interestingly, when the object is only shown *after* lifting, (i.e. object weight is perceived before size), no illusion is present ^16^, further supporting the view that the timing at which the size-weight illusion is formed might be critical.

The size and weight of objects are normally constant but can also be changed during object holding. For example, when pouring a drink into a transparent glass held in the hand, the size of the glass content as well as the weight of the filled glass change at the same time. It remains unclear whether *simultaneous* size and weight changes can mediate the SWI. To investigate whether size changes could induce illusionary weight changes, we simultaneously changed the size and weight of objects that participants held in the hand in a virtual reality environment and asked whether participants felt a weight increase or decrease. If the perceived weight change were affected by the size change in accordance with the SWI, a shrinking object (smaller) would lead to a perception of weight increase (heavier), whereas a growing object (larger) would induce an illusionary weight decrease (lighter). In a control experiment aiming at reproducing the classical SWI in our virtual reality setup, participant lifted large and small object sequentially as in standard SWI protocols ^3,5^. Since no expectation about object weight from its size can be formulated if they are perceived at the same time, we hypothesised that the SWI would be reduced when size and weight changes are perceived simultaneously after the object has been lifted from the table. This would indicate that the SWI indeed originates from an expectation about size-weight relationship and is not driven by bottom-up sensory information that is present throughout the time the object is manipulated.

## 2 RESULTS

We investigated the effect of simultaneous size and weight changes on the perception of weight change. We used a virtual reality environment that provided veridical visual information about size and object weight through haptic force feedback (Figure 1). Sixteen participants lifted an object by a yellow bar that was inserted into a blue cube of medium size. After the object was lifted, the blue cube could increase to a larger size (growing) or decrease to a smaller size (shrinking). Simultaneous to the change in size, the weight of the whole object could change as well. After replacing the object on the table, participants answered whether they perceived an increase or decrease in weight. A staircase procedure was used to determine the point where no perceptual weight change was perceived for the growing and shrinking conditions. Specifically, the staircase starts at a large increase or decrease in weight and adapts this change each trial until a weight change is no longer perceived. A psychometric curve was fitted to the answers (see Methods) to determine the perceptual bias. The difference between the point of no weight change perceived and the point where no actual weight change was introduced is the perceptual bias.

**Figure 1.**
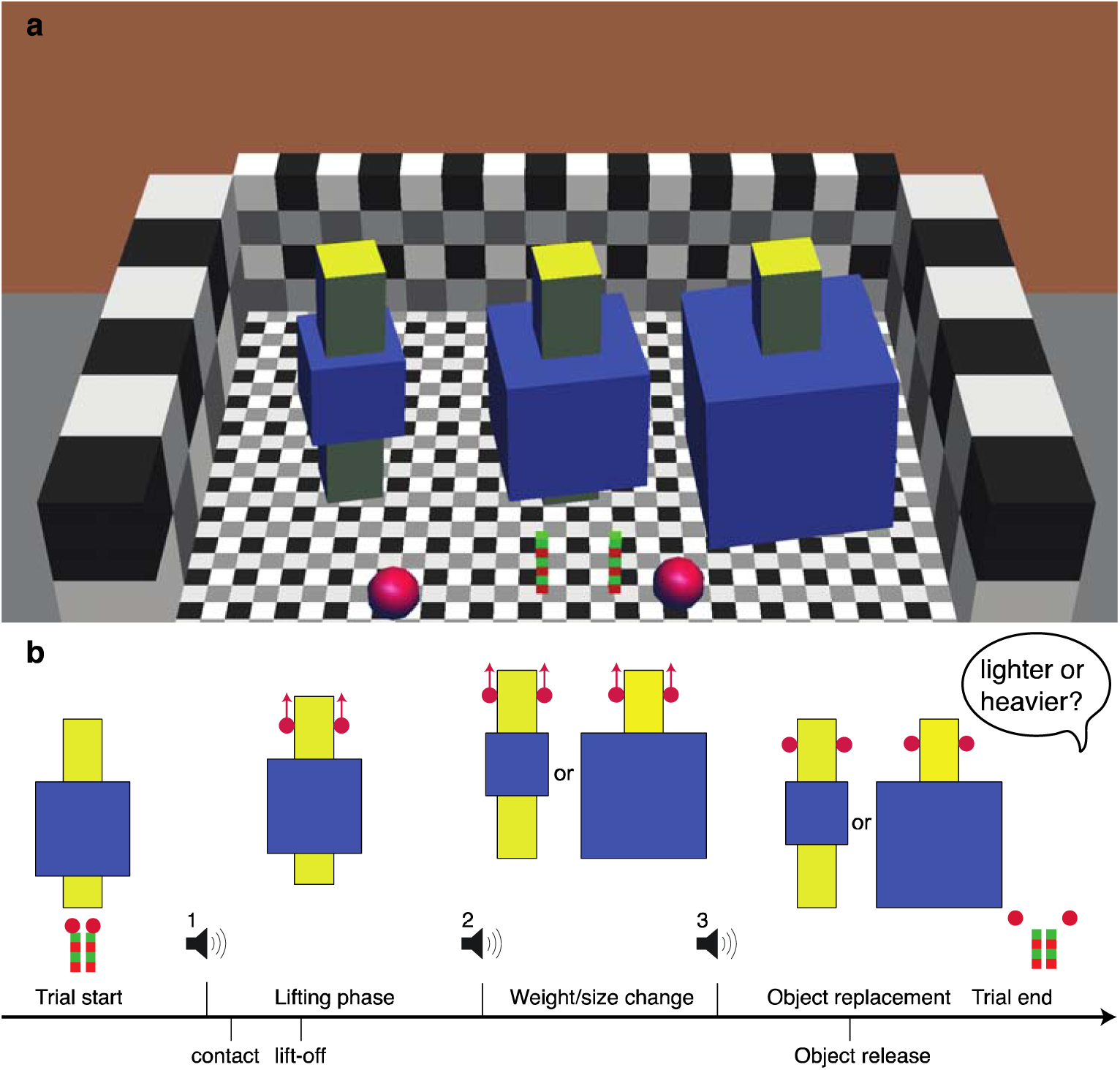
**a**: Virtual environment with lifted objects (yellow bar inserted into blue box), start positions (red-green poles) and fingers (red spheres). Small, medium (start) and large objects are shown. **b**: time line of the experiment. At the first beep, the medium object was lifted. Between the second and third beep, the object was changed, during holding, into a small or large object. After the change, the participant was asked to indicate whether the object became lighter or heavier.

Perceptual biases of weight change are shown in Figure 2, for shrinking and growing object separately. For the shrinking object, a bias of –0.52±2.8% was found, which was not significantly different from zero (t(15)=–0.18, p=0.86). When the object grew, a bias of 7.24±2.7% was found, which differed significantly from zero (t(15)=2.64, p=0.019). The total bias was the difference between the shrinking and growing conditions, divided by 2. The total bias was 3.88±1.3% and was significantly different from zero (t(15)=2.92, p=0.011). The psychometric fits also allowed us to determine discrimination thresholds of weight change perception. We divided the threshold by the initial weight before the change to calculate the Weber fraction. Weber fractions were 12.1±1.7% and 10.8±1.5% for shrinking and growing objects, respectively. Since Weber fractions indicate the discrimination threshold, this suggests that the biases that were found were smaller than the difference that can be reliably perceived.

**Figure 2.**
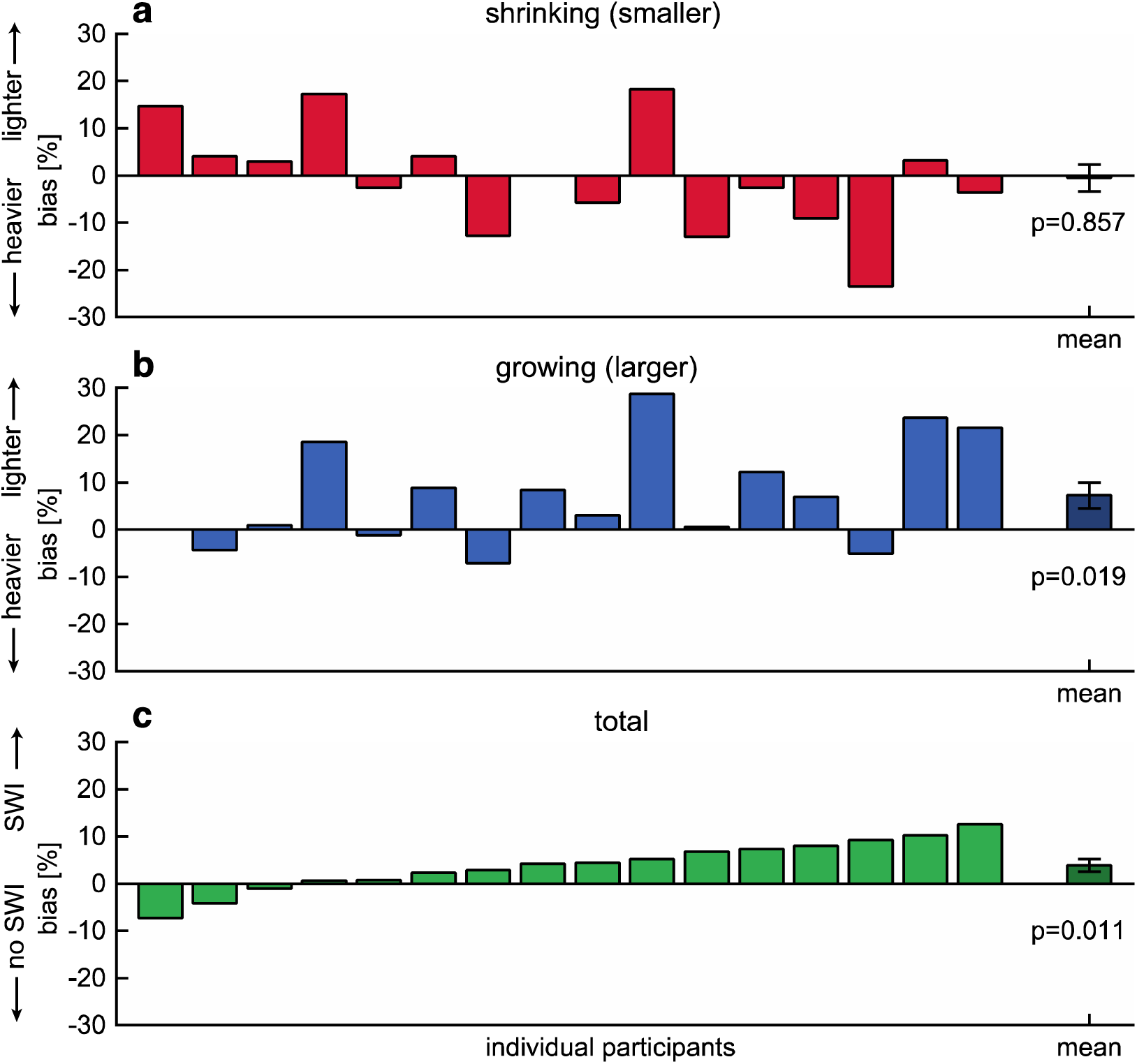
Biases for shrinking (**a**) and growing (**b**) objects, where positive or negative biases indicate the object is perceived to become lighter or heavier, respectively. **c:** Total biases, where positive biases indicate a size-weight illusion (SWI). Bars show values for individual participants. The rightmost dark bar represents the mean bias with standard error. p-values are shown for a one sample t-test, where growing and total biases were significantly different from zero.

In addition to perceptual measures, we also measured the forces applied by the participants and the position of the object. The changes in object height and grip force (GF) between the start and end of the change period are shown in Figure 3. As can be seen, object height varies with the object weight change, indicating that objects are held at a lower height when the weight increases and at a larger height when the weight decreases. When performing a linear regression on object height, slopes of −44.94 (95%CI:−49.9-40.0) and -40.12 (95%CI -43.9 -36.4) were found for shrinking and growing objects, respectively. When individual regression lines for shrinking and growing objects were compared with a paired samples t-test, no significant difference was found (t(15)=-1.71, p=0.109). This indicates that participants moved the object similarly in response to the weight change, regardless of the size change.

**Figure 3.**
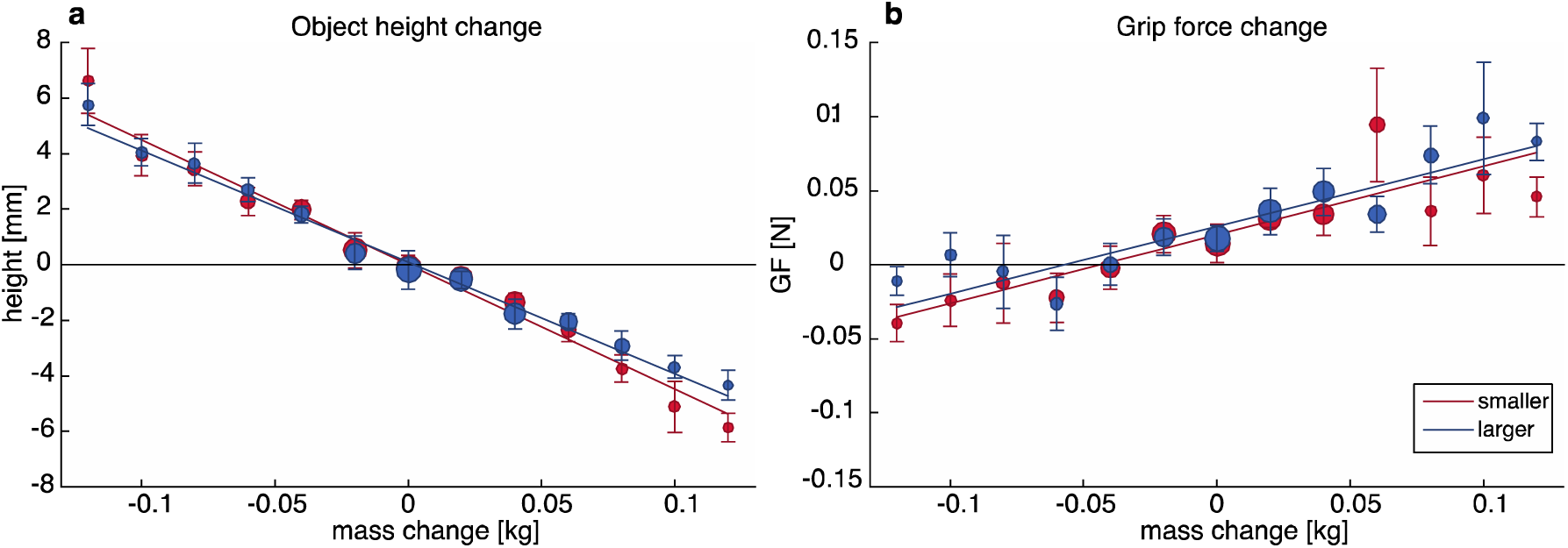
Changes in object height (**a**) and grip force (GF, **b**) during size/weight changes in static holding. Note that with increasing weight, the object moves down and grip force increases, similarly for smaller (shrinking, red) and larger (growing, blue) objects. Circle size represents the number of trials measured. Error bars indicate standard error.

In addition, GF increases with object weight change, indicating that more force is applied when the object becomes heavier and less force is applied when the object becomes lighter. For the linear regressions performed on GF, slopes of 0.46 (95%CI 0.28 0.64) and 0.45 (95%CI 0.30 0.60) were found for shrinking and growing objects, respectively. A paired t-test found no significant differences between individual regression lines for shrinking and growing objects (t(15)=0.134, p=0.895). This means that participants also adjusted their forces towards the weight change similarly for shrinking and growing objects.

### 2.1 Control Experiment

To ensure that we could reproduce the typical SWI in our virtual reality setup in a standard SWI paradigm, we performed a control experiment in which participants lifted small and large objects sequentially and directly compared them. Six participants who also participated in the main experiment, lifted the same small and large objects and responded which of the two was heavier. Again, we used a staircase procedure to determine the weight at which the two objects felt similar and fitted a psychometrical curve to the answers. Specifically, the weight of the test object was increased or decreased over the course of trials when it was perceived as lighter or heavier than the reference object, respectively. The difference in actual weight between the large and small object when they are perceived to be equal in weight reflects the perceptual bias. Multiple staircases were interleaved, where the small or the large object could be either the test or the reference object.

Biases in weight perception when comparing small and large objects are illustrated in Figure 4. On average, biases of –18.25±1.9% and 18.56±5.2% were found for a small test object and a large object test object, respectively. Both biases were significantly different from zero (small: t(5)=–9.43, p<0.001; large: t(5)=3.58, p=0.016). The total bias was calculated as the difference between the bias for the small and large object, divided by 2. A total bias of 18.41±3.5% was found, which was also significantly different from zero (t(5)=5.25, p=0.003). Weber fractions were 17.60±4.2% and 16.37±2.7% for the small and large test object, respectively.

**Figure 4.**
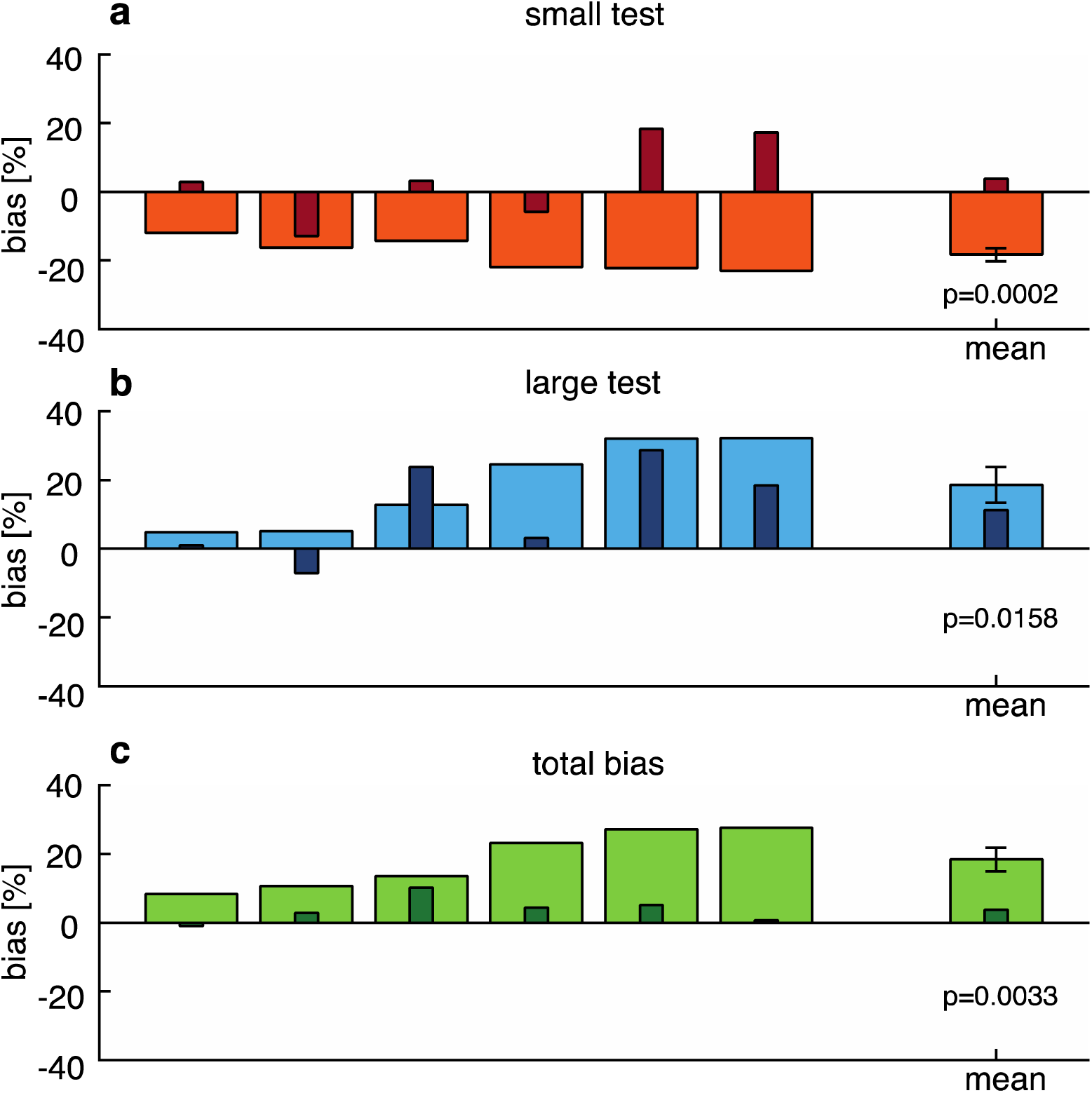
Biases for the control experiment (large bars) when comparing a large standard to a small test (**a**) or small standard with large test (**b**). Smaller, darker bars are the biases found in Experiment 1 for the same participants, for shrinking and growing objects, and total biases. Positive or negative biases indicate that the test object is perceived to be lighter or heavier than the standard, respectively. **c**: Total bias, with positive values indicating a size-weight illusion (SWI). Bars show biases for individual participants. The rightmost bar represents the mean with standard error. p-values are shown for a one sample t-test, where all biases were significantly different from zero.

To test whether small and large objects were lifted differently, we measured the forces participants applied when lifting the objects. Lifting parameters indicative of force scaling are illustrated in Figure 5 for the maximum grip force rate (GFRmax), maximum load force rate (LFRmax) and the loading phase duration (LPD). We compared the two object sizes for the reference weight (220 g) since we had the most trials for this weight. The lifting performance for small and large objects was comparable, as no differences were found in GFRmax (small: 8.6±1.2 N/s, large: 8.4±1.1 N/s, t(5)=0.76, p=0.479), LFRmax (small: 16.1±1.3 N/s, large: 16.5±1.4 N/s, t(5)=-0.64, p=0.551) or LPD (small: 0.27±0.03 s, large: 0.25±0.02 s, t(5)=1.67, p=0.157).

**Figure 5.**
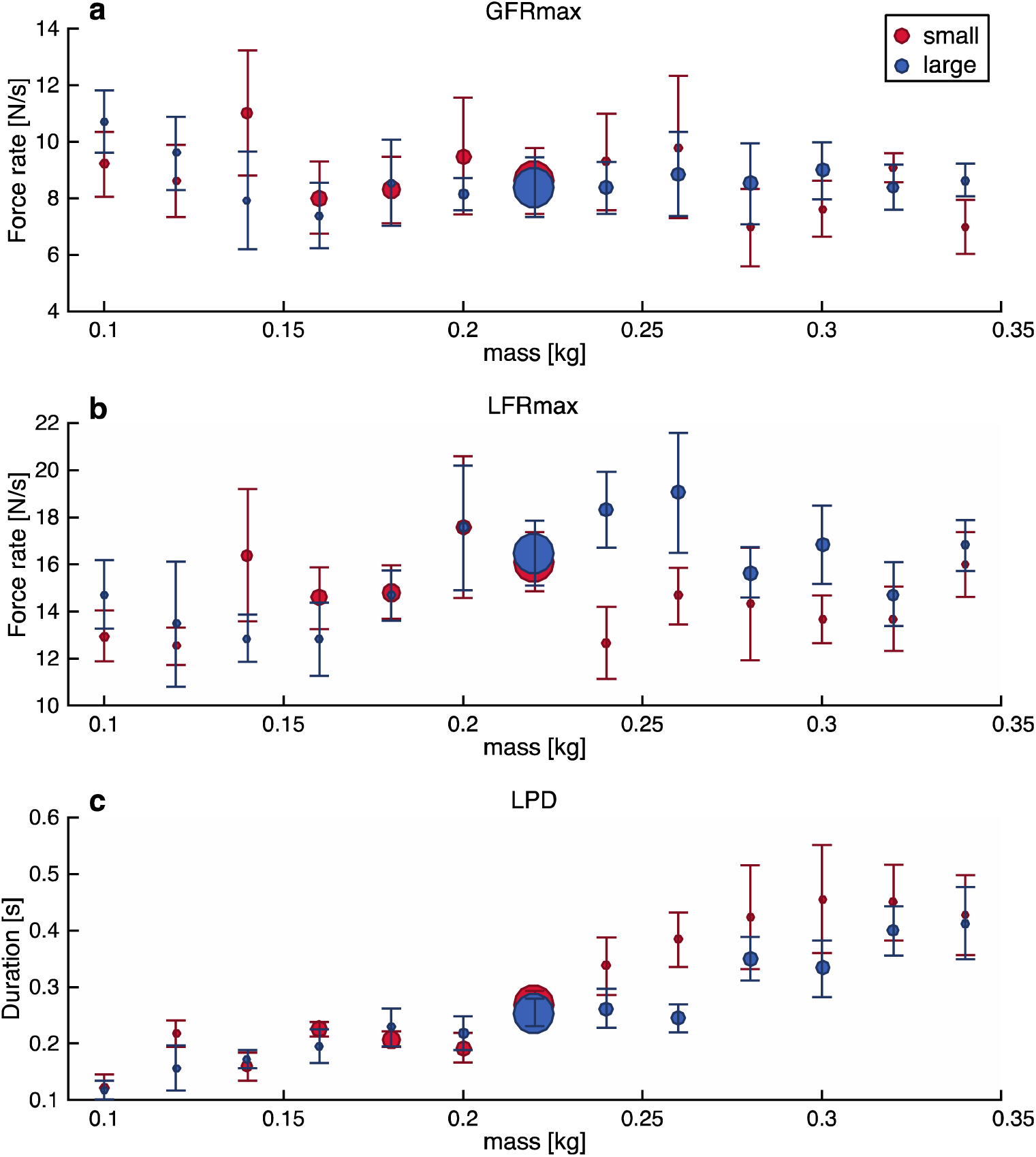
Peak grip force (GFRmax, **a**), peak load force (LFRmax, **b**) and load phase duration (LPD, **c**) for each lifted object weight. Note that force parameters are similar for small (red) and large (blue) objects. Circle size represents the number of trials measured. Error bars indicate standard error.

The biases found in the main experiment for the six participants who performed both experiments are also shown in Figure 4. The biases for the control experiment seem larger and more systematic than for the main experiment. This is also evident when comparing the average psychometric curves over all participants, as illustrated in Figure 6. If the biases from the control experiment were compared with the biases from the main experiment for the six subjects who performed both experiments, significant differences were found for a smaller object (t(5)=–3.60, p=0.016) and for the total bias (t(5)=–3.89, p=0.012). The difference was not significant for the large object (t(5)=1.58, p=0.174).

**Figure 6.**
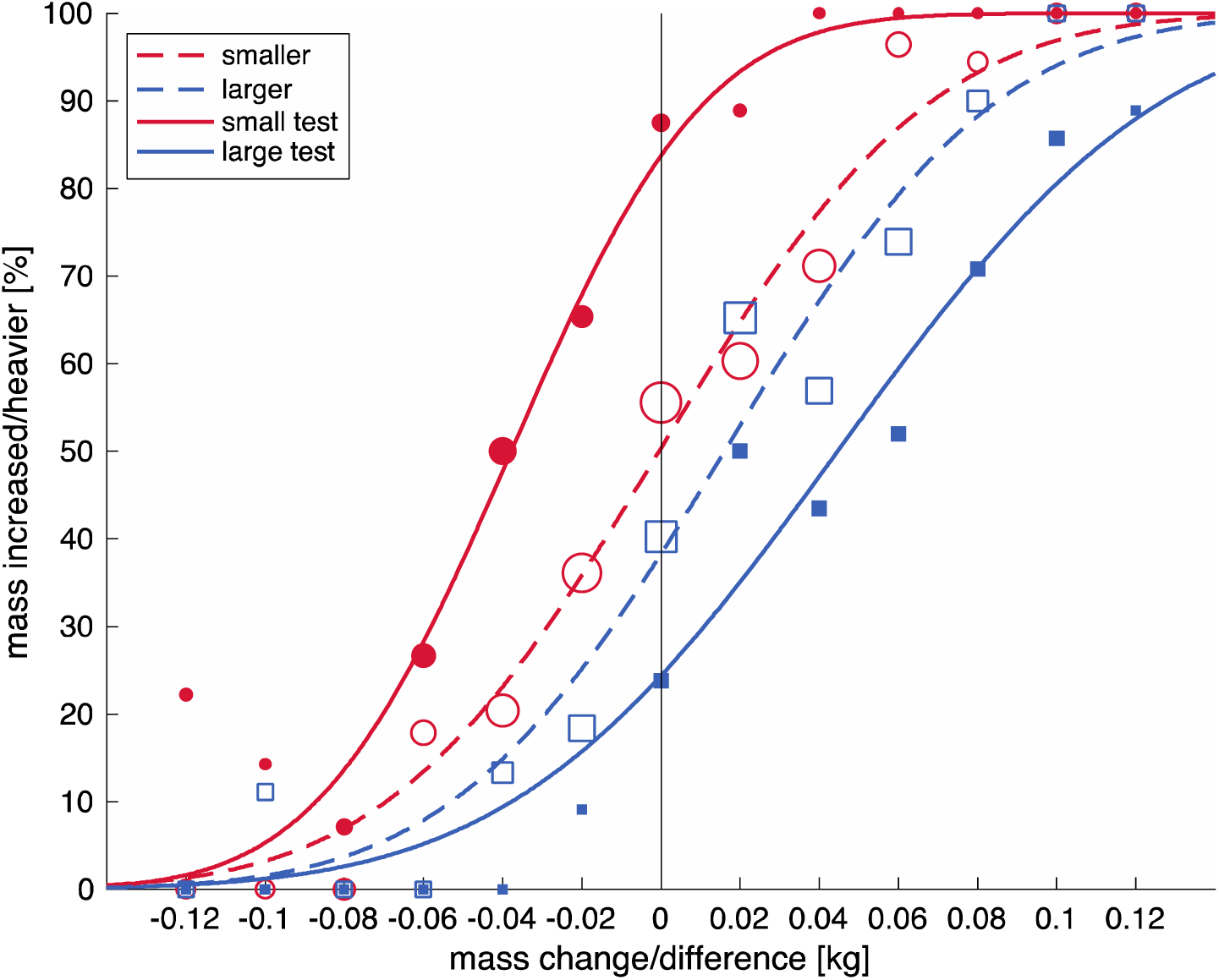
Psychometric curves for the main experiment (dashed lines, open symbols) and the control experiment (solid lines, close symbols), for small (red) and large (blue) objects. Symbol size represents the number of trials measured. Curves indicate the percentage where participants answered the test object to be heavier or the changed object to increase in weight for each lifted object weight or mass difference, respectively. Shifts to the left indicate objects feel heavier, shifts to the right indicate objects feel lighter. Note that the shifts are larger in the control experiment and in accordance with the size-weight illusion.

## 3 DISCUSSION

The aim of this study was to quantify the size-weight illusion (SWI) when size and weight were presented simultaneously after lifting the object. Secondly, the efficacy of simultaneous size-weight changes in mediating the SWI would also shed new light on the timing when SWI is formed. In the case of simultaneous size-weight changes after the object lift, no prediction of object weight can be derived from its size, we therefore hypothesized this would decrease the SWI, further accounting for the ‘prediction mismatch’ theory. Participants lifted virtual objects that changed in size and weight when they held the objects in the air. We found that changes in size induced perceptual biases of a smaller magnitude, where growing objects were perceived as decreasing in weight. On the other hand, no perceptual bias was found for shrinking objects. A control experiment demonstrated that the biases found in the main experiment were much smaller than when a standard SWI protocol was used in the same virtual reality setup. These results show that when object size and weight are presented at the same time, the SWI is substantially decreased.

The main results of the study showed that when the volume of an object was altered by a factor 2, a weight change of 4% was perceived. It must be noted that, whereas this total bias was significant, this effect is extremely small. We found that Weber fractions, indicating the relative difference that can be perceived, were much larger (11-12%) than the biases. Although the present experiment was not designed to measure accurate discrimination thresholds of weight change perception, these discrimination values are not very different from the literature on weight perception, where Weber fractions between 3-12% have been reported ^17^. The biases are smaller than weight differences that can be reliably perceived. This indicates that the shift in weight change perception induced by the size change falls within the range were perception is not very accurate. Therefore, the biases do not seem to be meaningful effects.

In contrast, the control experiment where small and large objects were lifted sequentially and compared to each other showed large biases that were larger than Weber fractions indicating a strong SWI. One can argue that the difference might be larger in this experiment because a small object and large object were directly compared, whereas in the main experiment the object changed from a medium size to a large or small object. However, even if the total bias of the mean experiment were doubled (8%), it is still smaller than the SWI (18%) we found in the control experiment.

Because of the difficulty to change the size and weight of objects simultaneously in various combinations with real-life objects, the present experiment was performed in a virtual reality environment. The control experiment allowed us to show that the SWI would also be present in our setup. It is noteworthy that the SWI has been shown in virtual environments before ^18–20^. Hence, the large reduction of the SWI when presented as a simultaneous size-weight change after lifting is not caused by the use of a virtual reality setup.

The perceptual effects that were found in our study do not seem to be related to differences in how the objects were handled. When objects increased in weight, the grip force was decreased and the object height decreased. Conversely, when objects decreased in weight, the grip force decreased and the object was moved slightly upwards. The adjustment in grip force is less pronounced with weight decreases. This might be because the safety margin is not compromised with a weight decrease, whereas it has to be increased with weight increases. Importantly, the same behaviour was seen for growing and shrinking objects. In addition, when lifting small and large objects of equal weight, force scaling was independent of object size. This is in agreement with earlier studies that showed independence of force control and weight perception with SWI objects ^5^.

Previous studies have suggested the SWI can be explained by bottom-up theories, such as considering weight perception as a combination of mass and density ^12,13^. However, this cannot explain the absence of an illusion in our experiment. If size and mass are presented simultaneously, the density of an object can be perceived at the same time as well and would still be expected to influence weight perception. The present results seem to be better explained by a top-down effect, where the expected weight based on object size is included into the final weight judgement ^2,6^. The perceived effect is then opposite to the initial expectation. In the SWI, larger objects are initially expected to be heavier, but then perceived as being lighter than small objects. With objects of changing size, a similar expectation can be formed that if an object grows, it would be expected to become heavier, but then feels as becoming lighter. Since in the present experiment the size and weight were changed simultaneously, no expectation of weight change based on size change would be formulated and could also not influence the perceived weight change. Nonetheless, since the change was slow, it is possible that expectations were still created but had a smaller impact. Moreover, the expectation might have had less effect because there is less time between the formation of an expectation and perceiving the actual consequence. Overall this led to a reduced SWI.

Our results showed no effect of shrinking objects on weight change perception, while growing objects did induce a significant bias where the object was perceived to become lighter. It is possible that this effect is partly influenced by a response bias towards lighter objects. Indeed, we saw that some participants show a bias shift in the same direction both for shrinking and growing objects. Perhaps these participants had a tendency to respond ‘lighter’ more often when they did not feel a weight difference. The decrease in weight might have been more difficult to perceive than the increase in weight. However, it is puzzling that such a response bias would only be apparent in the growing, not the shrinking condition. A more likely explanation would be that the growing object was more salient than the shrinking object. Especially since the yellow bar was present throughout the trial, which could have made the object appear long and therefore relatively larger in volume. However, our current virtual setup required the yellow bar to be present throughout the experiment to provide realistic interactions of the object with the environment and the fingers.

Furthermore, the timing at which expectations about object weight are incorporated into the perception might be very specific. The lifting phase might be a critical phase where expectations of weight are compared with sensory feedback. In a previous study we suggested that corrections to planned fingertip forces affected perception of object weight ^21^. Because in the current experiment size was altered only after lifting of the object had commenced, this suggest that comparisons in the lifting phase might be a critical time window for the SWI. The importance of this time period has also been noted in action observation, where weight must be inferred from observing the lifting movements of others ^22^. Moreover, a recent study showed that the SWI disappeared if size was shown after the object had been lifted ^23^. The importance of the lifting phase seems somewhat surprising, as the size-weight illusion is also found in situations where objects are not lifted, but placed or held in a supported hand ^9,24^, pushed when hanging from strings ^25^ or moved on a rail ^26^. However, placing an object in the hand or pushing it can also be considered as dynamic actions, which might be an important time period for evaluating expectations. Consistent with this notion, it has been found that the SWI is also seen if size information is shown during the replacement of a lifted object ^23^.

In conclusion, we found that that if size and weight are changed simultaneously after an object has been lifted, the size-weight illusion is greatly reduced. Since no prediction of weight based on size can be made in these conditions, this indicates that the size-weight illusion originates from a prediction mismatch. Furthermore, our findings suggest that the lifting phase might be a crucial time window for comparing weight expectations based on size with actual sensory information about object weight.

## 4 METHODS

### 4.1 Participants

Sixteen right-handed participants took part in the study (6 males, 10 females; age: 23±3.4, range 19-31 years). They all provided informed consent before the experiment and reported no visual deficits including lack of stereovision. All participants performed the main experiment. Six participants (3 males, 3 females, age: 23±2.2, range 20-26 years) also took part in the control experiment. The experiment was approved by the local ethical committee of KU Leuven.

### 4.2 Apparatus

We used a virtual environment to simulate objects that could be lifted and altered during holding. Two haptic devices (Phantom, Sensable) were placed underneath a mirror that reflected a 3D-screen (Zalman). The participants’ fingers could fit into the thimbles of the haptic devices and were unseen by the participant. In this way, a virtual environment was projected that was aligned with the actual positions of the participant’s fingers. The virtual environment (Figure 1) consisted of a patterned background to provide perspective cues, two start positions (red-green poles), the participants fingertips (red spheres) and the lifted object. We used a similar setup in a previous study ^27^.

The lifted object consisted of a yellow bar (12 cm height, 2.5 cm width) inserted into a blue cube. Before lifting the object, the blue cube was 6×6×6 cm and could be increased to a size of 8×8×8 cm or decreased to 4×4×4 cm. The yellow bar was not changed in size and the top part above the cube used for holding always had the same dimensions. The weight of the object always started at 220 g and could change with 120 g maximally, giving a weight range of 100-340 g for the final object weight. The weight was simulated for the complete object. The participants could interact with the yellow bar (i.e. it responded to virtual finger contact), whereas the blue cube was purely virtual.

The haptic devices provided force and position information in three directions, which were sampled with a 500 Hz frequency. Fingertip forces were determined in response to the position of the haptic devices, e.g. opposing forces were applied when the fingers contacted the virtual object. Normal forces applied to the cube (grip forces) were modelled as a spring, with an object stiffness of 400 N/m. Vertical (load) forces were the summation of the gravitational (g=9.81 m/s^2^), angular momentum and damping forces (damping constant is 2 kg/s). For the force calculations, the openHaptics toolkit was used embedded in custom-written software. See also ^27^ for further details on the set-up.

### 4.3 Task and procedure

Participants were seated in front of the virtual reality set-up and performed a few practice trials to get familiarized with the virtual environment and procedure. They were instructed to grasp and lift the object at the top part of the yellow bar. The time line of a trial is illustrated in Figure 1. Participants were instructed to coordinate their movements to three beeps. The first beep was played 1.5s after object appearance, the second beep at 4.5s and the third beep at 5.5s. Participants should commence their lift at the sound of a first beep, hold it in the air until two more beeps were heard and then replace the object on the table. In between the second and third beep, i.e. at a duration of 1s, the object would change in size and weight. After participants had replaced the object, they were asked to indicate whether the object had become heavier or lighter.

There were two size conditions, where the object could either grow to a large object, or shrink to a small object. The weight change was altered according to a staircase procedure. Four staircases were randomly interleaved: a shrinking object starting with a weight increase, a shrinking object starting with a weight decrease, a growing object starting with a weight increase and a growing object starting with a weight decrease. In this way, the order of increases and decreases in size and weight were randomly presented to the participant. The initial weight of the object was always 220 g. The staircases started at a weight change of 120 g (final weight 100 or 340 g) and were altered with steps of 20 g. For example, if a participant reported a weight increase of 120 g (340 g final weight) to become ‘heavier’, the next weight increase would be 100 g (320 g final weight). For each staircase, 15 trials were performed with a total of 60 trials. This was enough to for the staircases starting at weight increases and decreases to converge, which was confirmed by visual inspection of the data.

### 4.4 Control experiment

Six participants also performed a control experiment of a standard size-weight illusion task. They sequentially lifted two objects, a small and a large one, and responded which of the two was heavier. A staircase procedure was used to determine the weight combination at which the small and large object felt equal. Four staircases were randomly interleaved: small standard starting with a large light test weight, small standard starting with large heavy test weight, large standard starting with a small light test weight, large standard starting with a small heavy test weight. The standard object had a weight of 220 g, which was equal to the initial weight in the main experiment.

The weight of the test object in the following trial was adapted based on the answer of the participant. The start weights of the test objects were light (100 g) or heavy (340 g). The weight of the test object was increased with 20 g if the participant reported the test object to be lighter or decreased with 20 g if the test object was perceived as being heavier. The presentation order of the small and large object was randomized across trials. For each staircase, 15 trials were performed with a total of 60 trials. This was enough for staircases to converge, which was confirmed by visual inspection of the data.

### 4.5 Perceptual analysis

For the main experiment, the percentage of trials where participants reported the object as becoming ‘heavier’ was calculated for each size change and each weight change. Through these percentages a weighted psychometrical curve was fitted to determine the bias (µ) and standard deviation (*σ*):

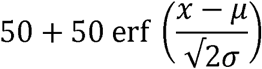

The bias was determined for shrinking and growing objects separately, where a positive bias indicates that objects are perceived as being lighter. The biases were expressed as a percentage change with respect to the initial weight (220 g). The total bias was calculated as the bias for the shrinking object subtracted from the bias for a growing object and dividing this number by 2.

For the control experiment, the percentage where the test object was perceived as being ‘heavier’ was calculated for each weight difference between the object pairs. This was done separately for conditions with a small and with a large test object. A psychometrical curve was fitted through these percentages and the bias was expressed as a percentage with respect to the standard weight (220 g). The total bias (size-weight illusion) was calculated as the bias for the small test object subtracted from the bias for a large test object and dividing this number by 2.

Discrimination thresholds were determined from *σ*, which corresponds to the difference between 50% and 84% correct answers of weight change. We expressed the threshold as a percentage of the initial weight (or standard weight in the control experiment) to calculate the Weber fraction.

### 4.6 Force and position analysis

To see whether there were any differences in behaviour in response to shrinking or growing objects (main experiment) or between lifting small or large objects (control experiment, we also measured fingertip forces and object position in both experiments. Force and positional data were filtered with a bidirectional 4^th^ order low-pass Butterworth filter, with a cut-of frequency of 15 Hz. Missing samples of the time series (<0.02%) were linearly interpolated. Grip force (GF) was determined as the normal force applied perpendicular to the object, averaged over both fingers. Load force (LF) was determined as the vertical forces tangential to the object, summed for both fingers. Object height was the vertical distance from the virtual table to the centre of the object.

For the main experiment, only parameters during the change of the object weight during holding were considered. The change in object height and GF were computed for each size and weight change by subtracting the values at the onset and end of the change period (i.e. values at the third and second beep).

For the control experiment, we looked at parameters indicative of motor planning, which are the force rates and the load phase duration (LPD). The force rates were the time derivatives of the forces. The variables of interest were the peak force rates (GFRmax and LFRmax), which were determined between 50 ms before GF onset and 50 ms after lift-off. Force onset was set at the first point that the force reached a value of 0.1 N and continued to a value of at least 1N. Lift-off was the time point at which LF overcame object weight. The LPD was the time between LF onset and lift-off.

### 4.7 Statistical analysis

One sample t-tests were used to determine whether perceptual biases were significantly different from zero. To compare the biases between the main and control experiment, the data of the participants who performed both experiments (N=6) were compared with paired-samples t-tests.

For the main experiment, mean changes in object height and GF were plotted against object weight change. Because not all weight changes were presented equally, a weighted linear regression was performed on these parameters, with the number of trials as weights.

For the control experiment, only the force parameters for lifting an object with a standard mass (220 g) were statistically compared, because too few trials were performed with other object weights. Paired samples t-tests were performed on GFRmax, LFRmax and LPD to test differences between the lifts of small and large objects.

Due to technical errors, 13 trials were removed from the perceptual analysis (1%) in the main experiment. In addition, trials in which the object was lifted multiple times, lifted just after initiation of the change or dropped before the end of the change period were removed from the force analysis as well (total 63 trials, 7%). In the control experiment, trials in which objects were lifted multiple times were removed from the force analysis (13 trials, 2%).

## ACKNOWLEDGMENTS

This research was supported by Fonds Wetenschappelijk Onderzoek grants to VVP (FWO post-doctoral fellowship, Belgium, 12X7118N) and MD (FWO Odysseus, Belgium, G/0C51/13N).

## AUTHOR CONTRIBUTIONS

VVP and MD designed the experiments. VVP collected and analysed the data. VVP wrote the intial version of the manuscript. VVP and MD both reviewed the manuscript.

## DATA AVAILABILITY

The authors can make the data available upon request.

